# Histological analysis of implantation embryos in large Japanese field mouse (Apodemus speciosus) and estimation of developmental stage

**DOI:** 10.1101/2021.03.24.436754

**Authors:** Hiroyuki Imai, Kiyoshi Kano, Ken Takeshi Kusakabe

## Abstract

The large Japanese field mice (*Apodemus speciosus*) are small rodent specie endemic to Japan. The genetic characteristics of *A. speciosus* is different chromosome numbers within one species. Furthermore, *A. speciosus* is used for research in radiation and genetics. In this present study, a pregnant *A. speciosus* was obtained, and histochemical analysis of the implanted embryos was performed and compared with developmental stages of the mouse. Although there were some differences, the structures of the implanted embryos including the primitive streak and placenta of *A. speciosus* were similar to that of the mouse. Our study will be important report in the construction of a developmental atlas of *A. speciosus*.

## Main Text

The large Japanese field mice (*Apodemus speciosus*) is a small rodent that inhabits Japan and have diverged from its mouse and rat ancestor 1.5-4.4 hundred thousand years ago [3]. *A. speciosus*, found throughout the Japan except for Okinawa, is used as a model for studying geographic isolation of animals [11]. In addition, *A. speciosus* has distinctive genetic characteristics in that the number of chromosomes differs between western and eastern Japan. This karyotypic alternation in *A. speciosus* was caused by a Robertsonian translocation [10]. In recent years, *A. speciosus* has been used as animals to monitor the effects of radiation from nuclear power plant accidents [1].

During the genetic analysis of *A. speciosus*, which is our current project, A pregnant mice was obtained by chance. The morphology of the implantation sites was compared with Theiler’s house mouse developmental staging [12] and summarized as a case report.

With the permission, *A. speciosus* were captured using the sharman traps (LFA; Hoga, Kyoto, Japan). Each implantation site was identified (Fig. S1), fixed with Bouin solution and routinely embedded in paraffin. Serial section of 5μm were prepared using RM2245 (Leica, Wetzlar, German) and stained with H.&E.

The observation revealed typical primitive streak-like structures and two-three cavities except for R1 conception (Fig. 1). The primitive streak-like structures were similar to that of mouse embryos and consisted of three layers of cells, suggesting that the primitive streak structures containing these three cell layers were the primitive streak of *A. speciosus* (Fig. 1 Diagram; Pr). There is a region that binds the maternal decidua and conception on the opposite side of the primitive streak, which is assumed to be the placental region (Fig. 1 Diagram; Pl). Based on the orientation of primitive streak and placenta, the three cavities in the conception appeared to be ectoplacemtal cavity, exocoelom and amniotic cavity, respectively (Fig. 1 Diagram; Ec, Ex, A). Even in L1 preparation, where only two cavities were founded, part of partition between ectoplacental cavity and exocoelom could be observed. For R1 preparation, where only one cavity was founded, this might be a transverse section of primitive streak region, as there were three layers of cellular structures. Thus, these results indicated that the mice embryos used in this present study was 7.5-8.0 days old, which is corresponded to stage 11. However, there were some features that do not completely correspond the Theiler’s developmental stages of mouse embryo; 1. allantois, which is stage-specific structure in exocoelom, could not be observed, 2. no clear blood islet could be observed. Therefore, *A. speciosus* embryos shows morphology basically similar to that of mouse development, but they also might differ slightly from mouse development.

**Figure 1.**
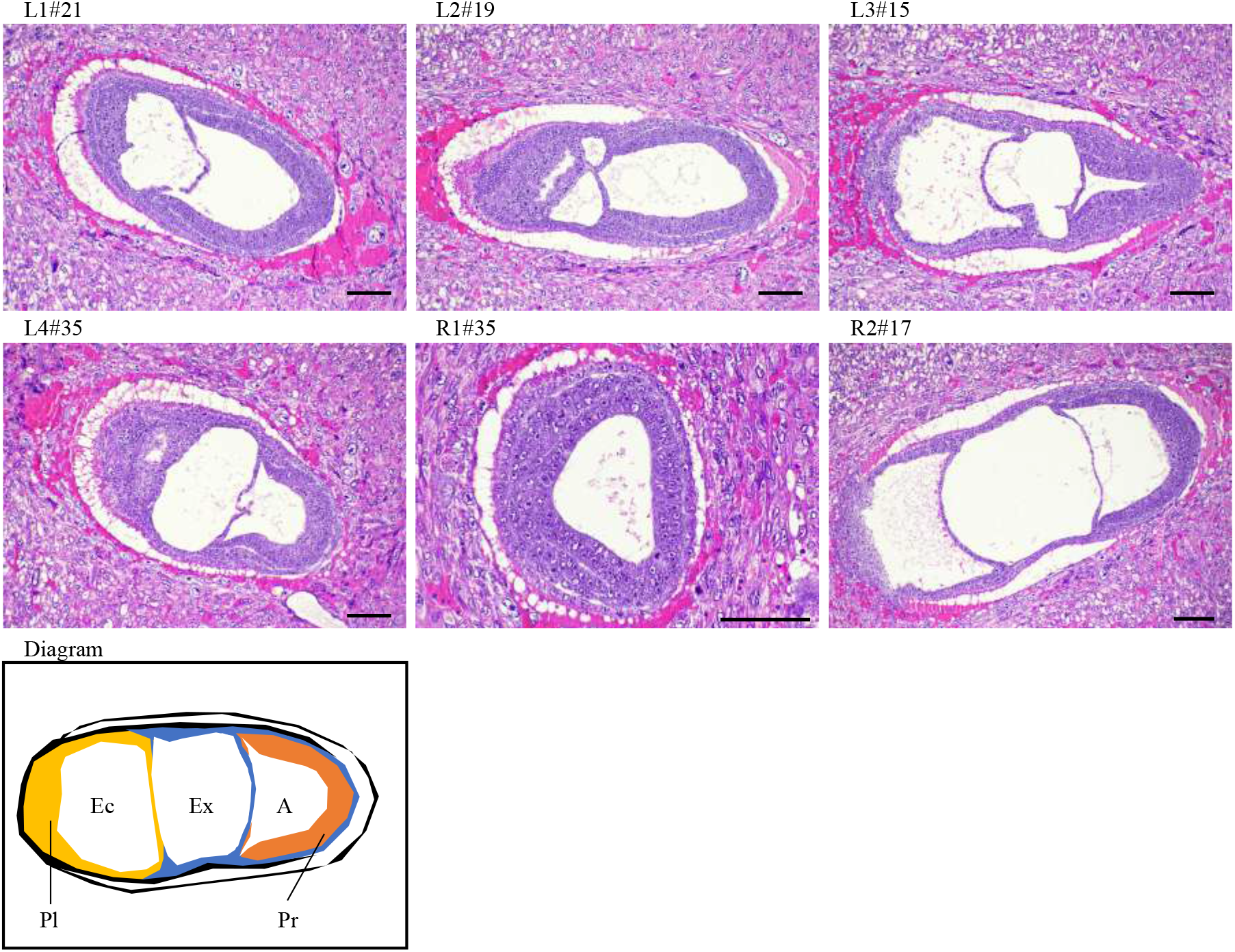
Histological images of implantation sites in the Apodemus speciosus. The implantation site number (L1-R2) and the preparation number of the serial sections (#__) were shown in the upper left of images. The abbreviations in the diagram were as follows; Pl: Placenta, Ec: Ectoplacemtal cavity, Ex: Exocoelom, A: Amniotic cavity and Pr: Primitive streak. Bar; 100 μm.

As above, this present study showed the embryonic morphology of the stage 11 of *A. speciosus*. Although these embryonic ages were unknown, the gestation period of *A. speciosus* was 21-26 days [7], these embryos might be 7.9-10.4 days old. Methods of rearing and breeding the *A. speciosus* were being developed [9], and an atlas of embryonic development of *A. speciosus* could be created in the future. Furthermore, the whole genome sequence of *A. speciosus* had already been published [5], and *A. speciosus* can be established as a novel experimental rodent with the following characteristics; 1. rodents with seasonality in spermatogenesis and reproduction [4, 8], 2. with different chromosome numbers within the same species, 3. pioneering developmental engineering [2, 6]. This present study will be important as a morphological report of on point in the developmental atlas of *A. speciosus*.

## Acknowledgements

This study was supported by Research and survey program of Akiyoshidai Karst Plateau Academic Center, Yamaguchi University to I.H. The authors thanks Laboratory of Veterinary Anatomy and Embryology, Yamaguchi University for participation in animal trapping.

---

**Figure S1.**
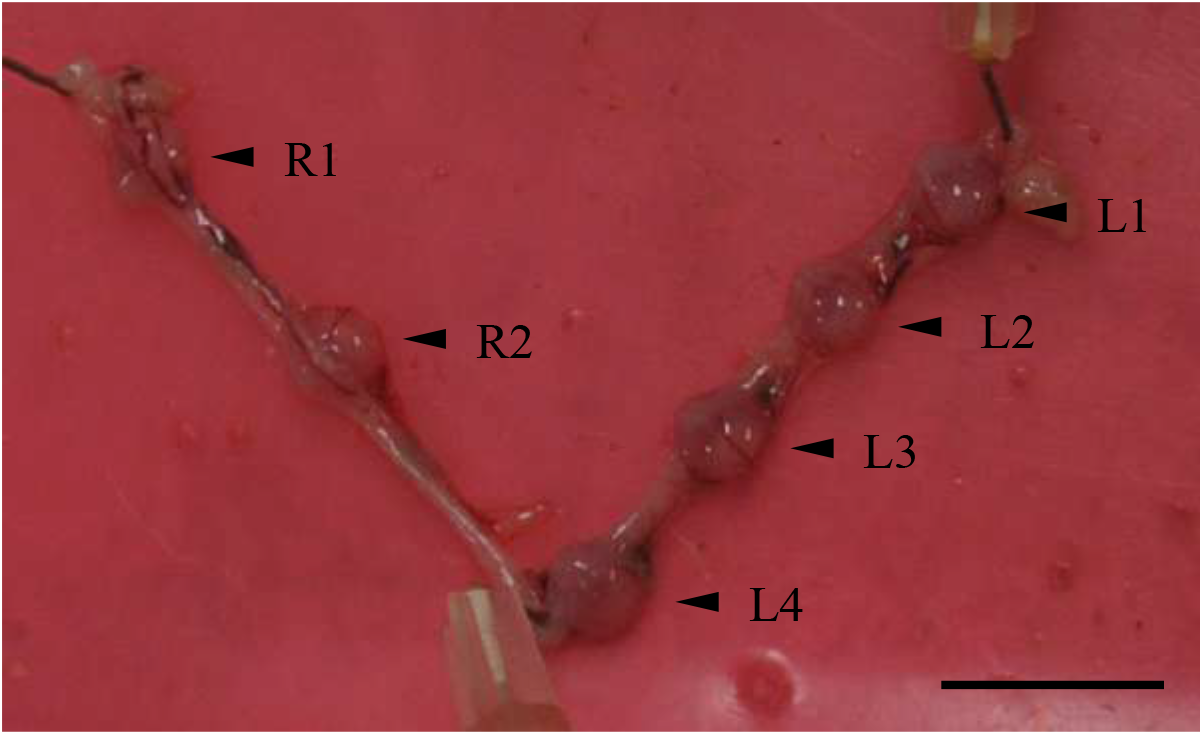
The implantation uterus of Apodemus speciosus. The arrowheads indicated the implantation site number (L1-R2). Bar; 10 mm.

## References

1. Ariyoshi, K., Miura, T., Kasai, K., Akifumi, N., Fujishima, Y. and Yoshida, M.A. 2018. Radiation-induced bystander effect in large Japanese field mouse (Apodemus speciosus) embryonic cells. Radiat. Environ. Biophys. 57: 223–231.

2. Azuma, R., Hatanaka, Y., Shin, S.W., Murai, H., Miyashita, M., Anzai, M. and Matsumoto, K. 2020. Developmental competence of interspecies cloned embryos produced using cell from large Japanese field mice (Apodemus speciosus) and oocyte from laboratory mice (Mus musculus domesticus). J. Reprod. Dev. 66: 255–263.

3. Hanazaki, K., Tomozawa, M., Suzuki, Y., Kinoshita, G., Yamamoto, M., Irino, T. and Suzuki, H. 2017. Estimation of evolution rates of mitochondrial DNA in two Japanese field mouse species based on calibrations with quaternary environmental changes. Zoolog. Sci. 34: 201–210.

4. Kuwahara, S., Mizukami, T., Omura, M., Hagihara, M., Iinuma, Y., Shimizu, Y., Tamada, H., Tsukamoto, Y., Nishida, T. and Sasaki, F. 2000. Seasonal changes in the hypothalamo-pituitary-testes axis of the Japanese wood mouse (Apodemus speciosus). Anat. Rec. 260: 366–372.

5. Matsunami, M., Endo, D., Saitou, N., Suzuki, H. and Onuma, M. 2017. Draft genome sequence of Japanese wood mouse, Apodemus speciosus. Data Brief 16: 43–46.

6. Meguro, K., Komatsu, K., Ohdaira, T., Nakagata, N., Nakata, A., Fukumoto, M., Miura, T. and Yamashiro, H. 2019. Induction of superovulation using inhibin antiserum and competence of embryo development in wild large Japanese field mice (Apodemus speciosus). Reprod. Domest. Anim. 54: 1637–1642.

7. Murakami, O. 1974. Growth and development of the Japanese wood mouse (Apodemus speciosus) I. The breeding season in the field. Japanese Journal of Ecology. 24: 194–206. (In Japanese, Abstract in English)

8. Okano, T., Onuma, M., Ishikawa, H., Azuma, N., Tamaoki, M., Nakajima, N., Shindo, J. and Yokohata, Y. 2015. Classification of the spermatogenic cycle, seasonal changes of seminiferous tubule morphology and estimation of the breeding season of the large Japanese field mouse (Apodemus Speciosus) in Toyama and Aomori Prefectures, Japan. J. Vet. Med. Sci. 77: 799–807.

9. Sakai, Y., Sakamoto, S.H., Kato, G.A., Iwamoto, N., Ozaki, R., Eto, T., Shinohara, A., Morita, T., and Koshimoto, C. 2013. Rearing method to induce natural mating of the large Japanese field mouse, Apodemus speciosus. Honyuruikagaku 53: 57–65. (In Japanese, Abstract in English)

10. Shimba. H, and Kobayashi, T. 1969. A Robertsonian type polymorphism of the chromosomes in the field mouse, Apodemus speciosus. Jpn. J. Genet. 44: 117–122.

11. Suzuki, Y., Tomozawa, M., Koizumi, Y., Tsuchiya, K. and Suzuki, H. 2015. Estimating the molecular evolution rates of mitochondrial genes referring to quaternary ice age events with inferred population expansion and dispersals in Japanese Apodemus. BMC Evol. Biol. 15: 187.

12. Theiler, K. 1989. The house mouse: Atlas of embryonic development. Springer-Verlag. New York.

